## Supplementary material for "The Rapidly Evolving X-linked *miR-506* Family Finetunes Spermatogenesis to Enhance Sperm Competition": fig. S

**This PDF file includes:**

Figures S1 to S9

**Other supporting materials for this manuscript include the following:**

Tables S1 to S8

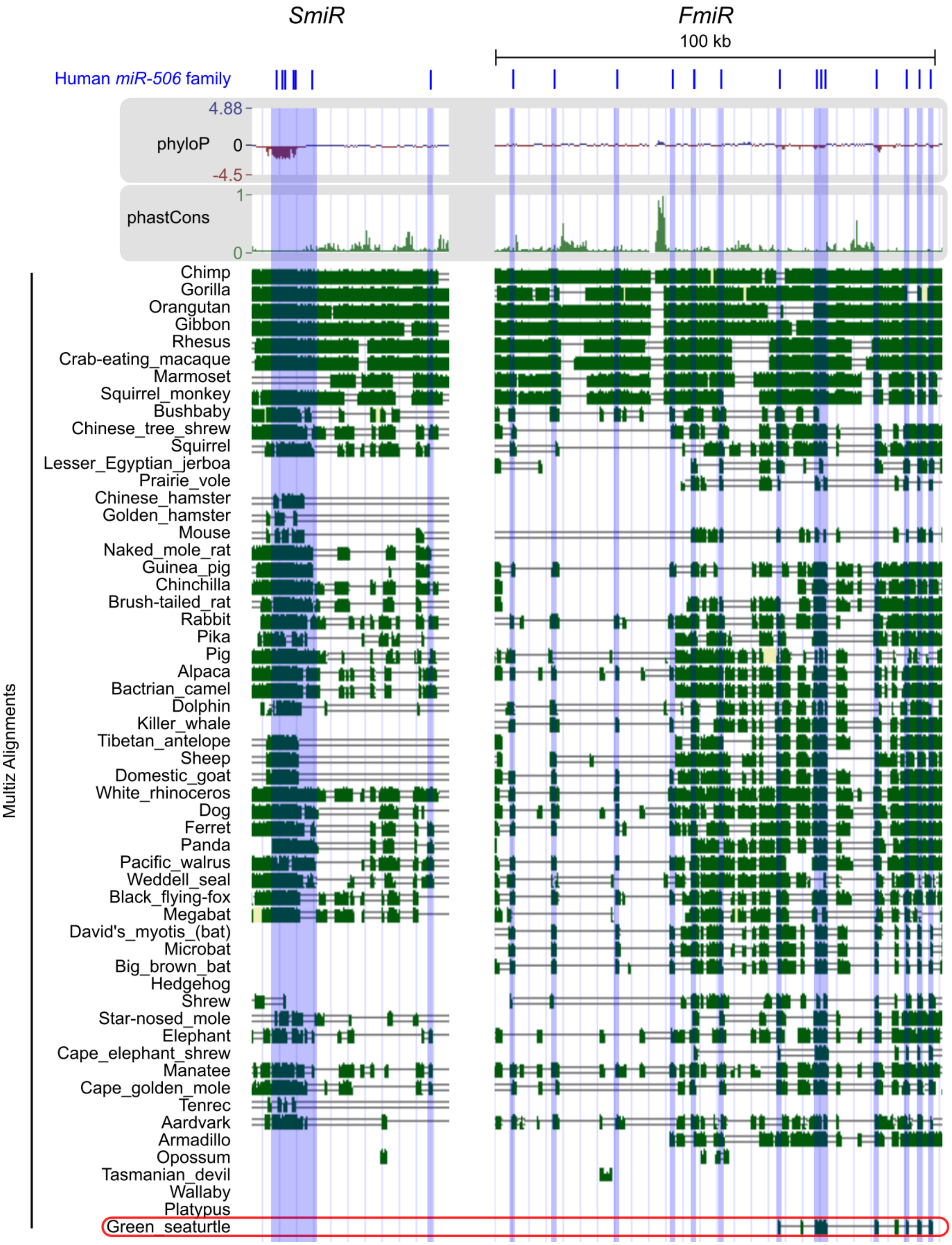


**Figure S1. Multiz Alignment & Conservation analyses of X-linked *miR-506* family using the human genomes as references.** Multiz Alignment & Conservation analyses of the X-linked *miR-506* family across 100 species using the human genome as the reference. miRNAs are highlighted in blue.

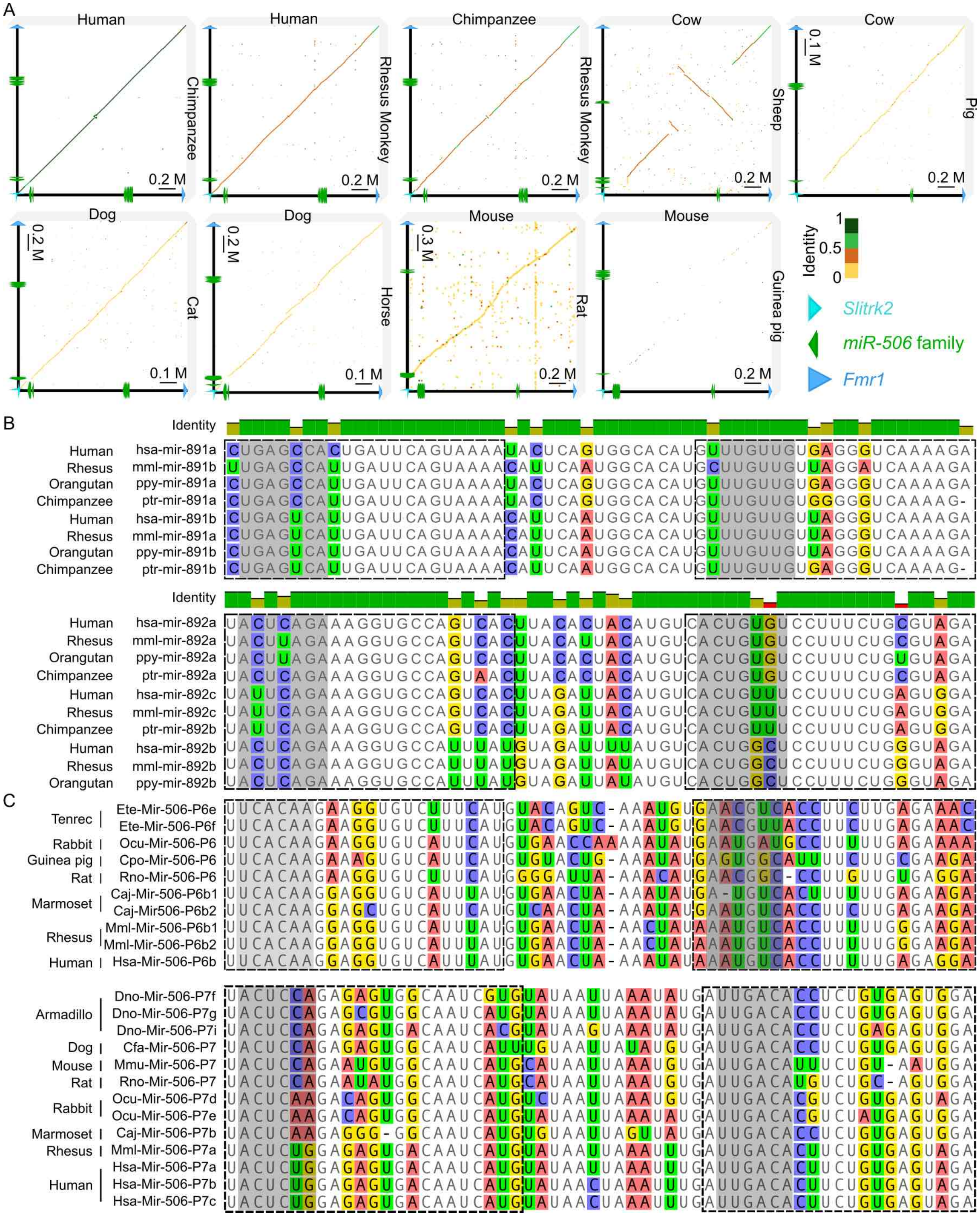


**Figure S2. Genomic and sequence similarity among members of the X-linked *miR-506* family.** A. Genomic similarity using GENIES-based dot plot analyses within clades. B. Upper panel, sequences alignment of precursor *miR-891* among primates. Lower panel, sequences alignment of precursor *miR-892* among primates. Bases in the dashed square represent the mature miRNA sequences. Seed sequences are highlighted in grey, and alignments with mismatches are indicated with other colors. C. Alignment of precursor miRNA of *miR-506-P6* (upper panel) and *miR-506-P7* (lower panel) subfamily across species. Bases in the dotted square represent the mature miRNA. Seed sequences are shown in the grey background, and alignments with mismatches are indicated in bases with the colored background.

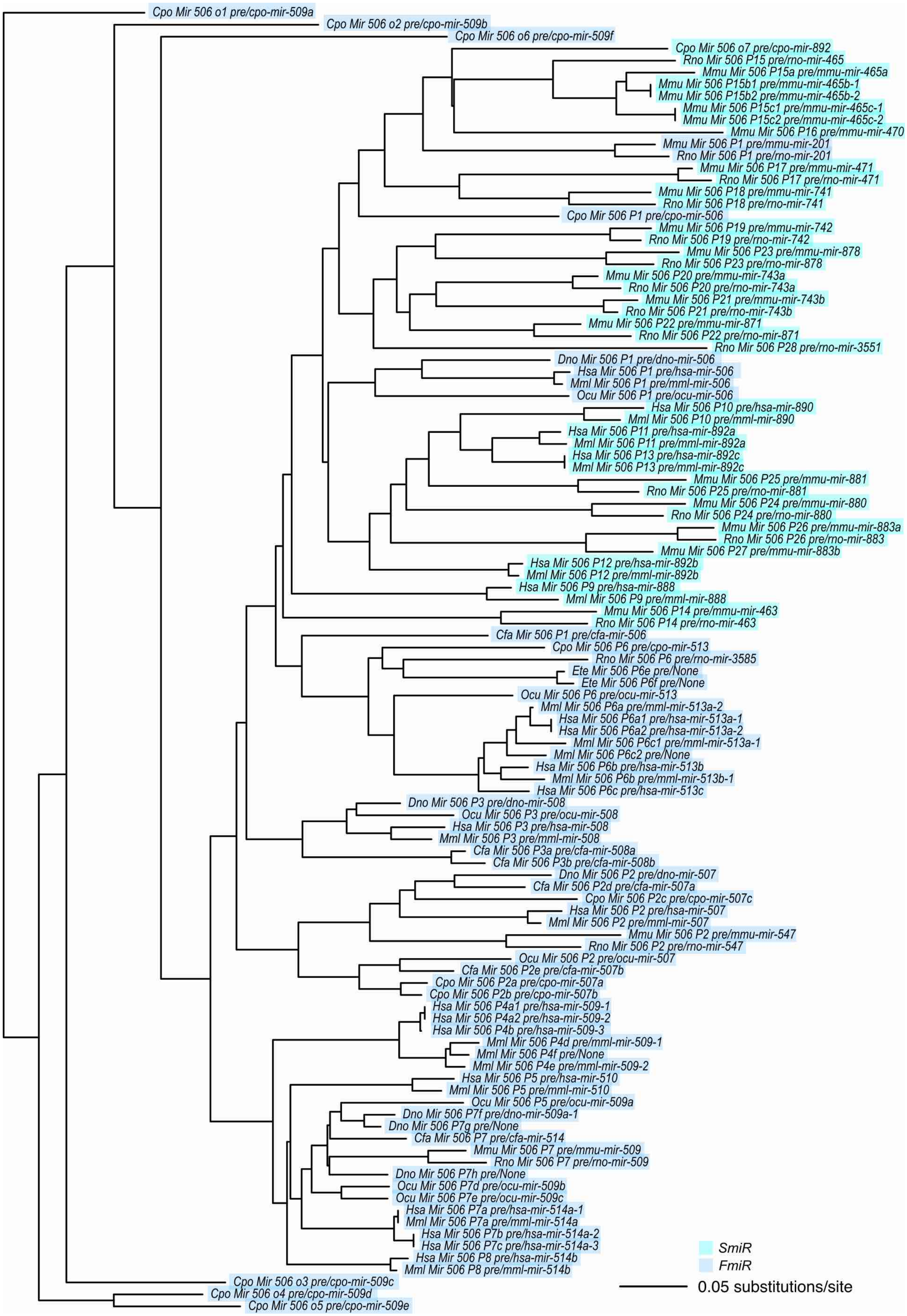


**Figure S3. A phylogram of the X-linked *miR-506* family.** SmiR and FmiR are shown in cyan and blue, respectively.

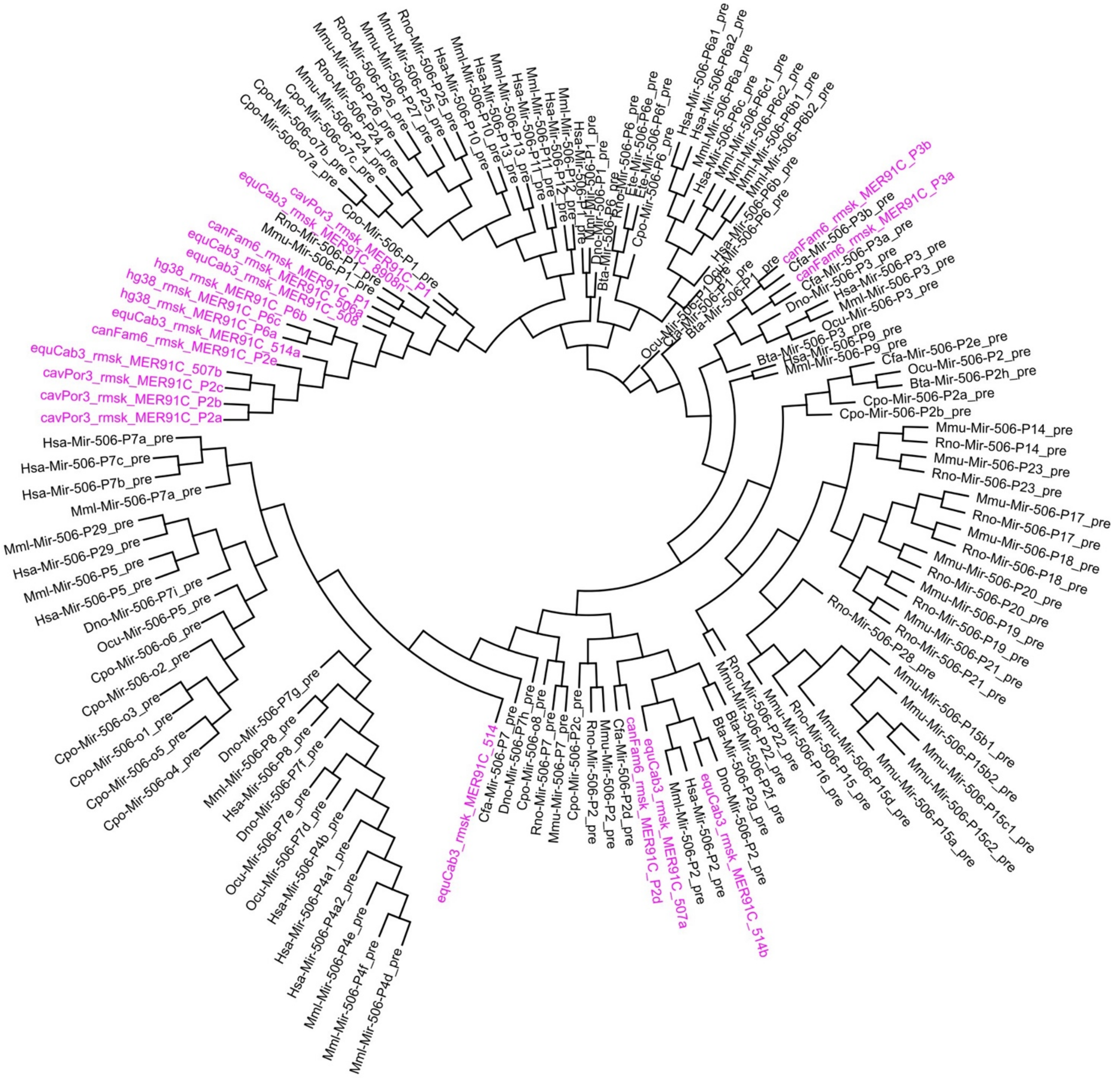


**Figure S4. A phylogenetic tree of the MER91C DNA transposons and the X-linked *miR-506* family miRNAs.** The MER91C DNA transposons are labeled in purple.

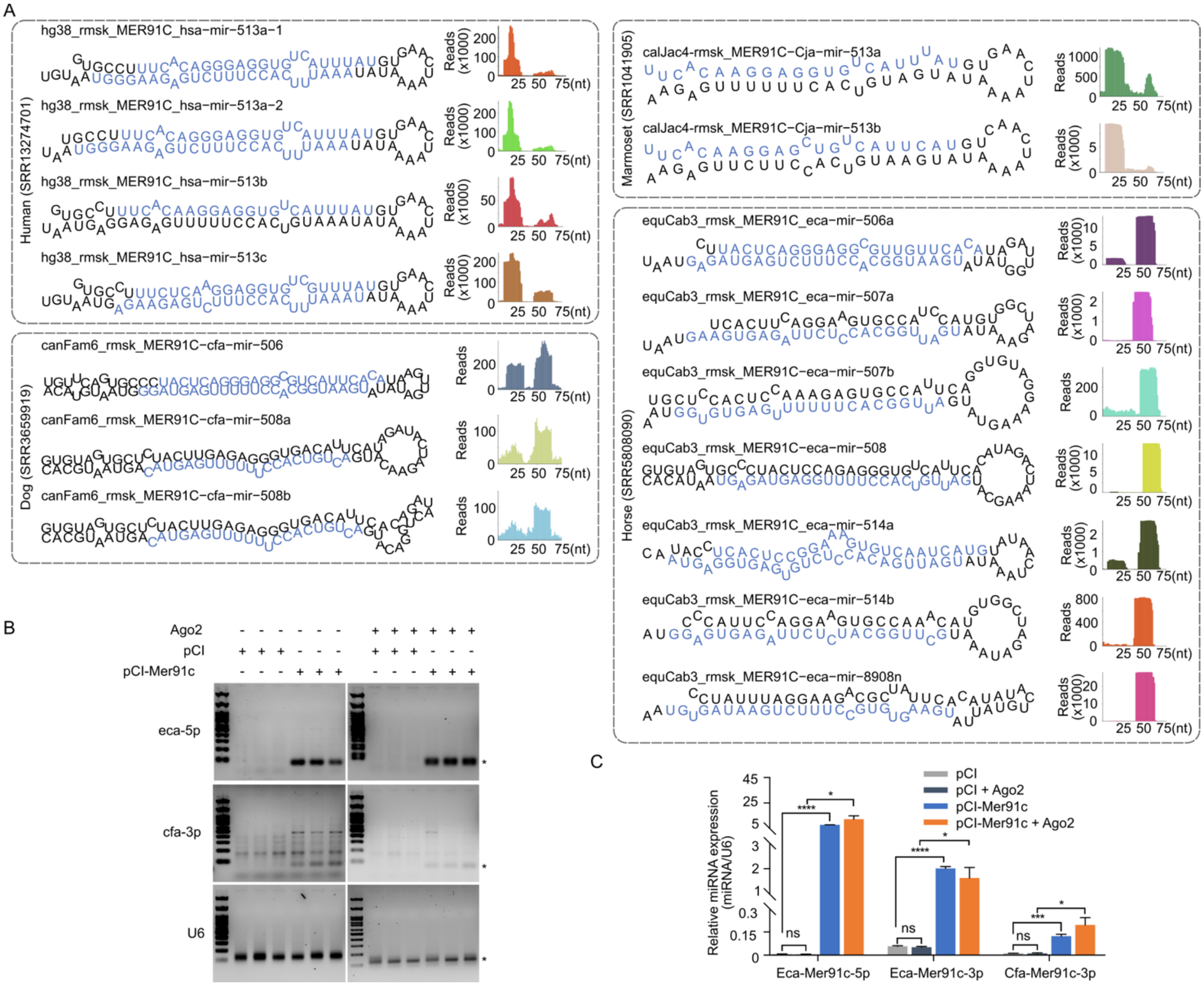


**Figure S5. X-linked *miR-506* family is derived from MER91C DNA transposon and expanded *via* LINE retrotransposons.**

A**.** RNA structure and sRNA-seq reads of the MER91C DNA transposon-derived miRNAs in humans, marmosets, horses, and dogs. The mature miRNAs are labeled in blue in the RNA structure. B. A representative gel image of the MER91C DNA transposon-derived miRNAs from horses and dogs expressed in HEK293T cells. n=3 for each sample. * indicates the expected size of the miRNA. U6 was used as the internal control. C. qPCR of the MER91C DNA transposon-derived miRNAs expression in HEK293T cells. n=3 for each sample. *, ***, **** indicate $p < 0.05$, 0.001, 0.0001, respectively. One-way ANOVA was used for the statistical analysis.

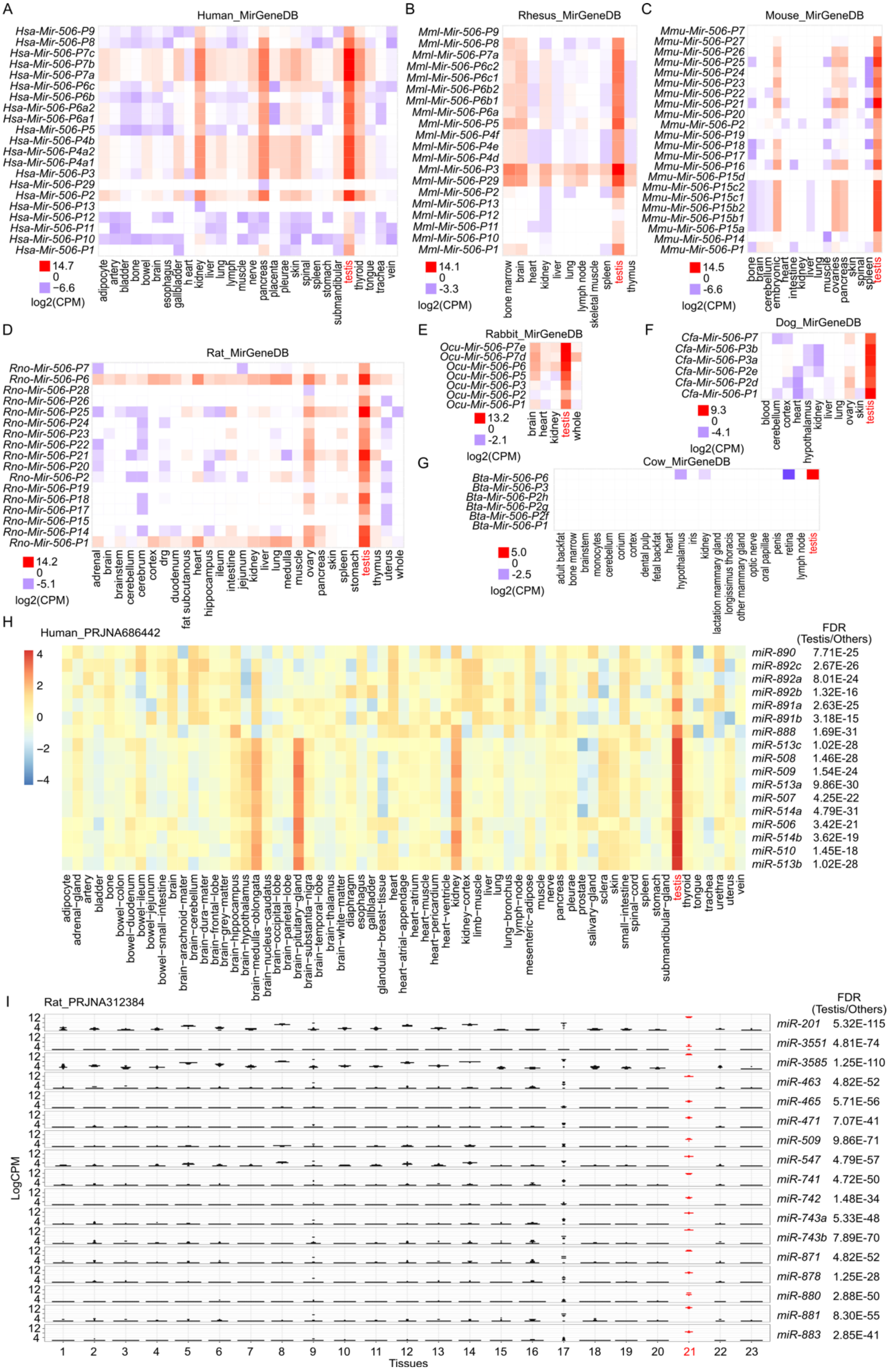
A
Human_MirGeneDB
Hsa-Mir-506-P9
Hsa-Mir-506-P8
Hsa-Mir-506-P7c
Hsa-Mir-506-P7b
Hsa-Mir-506-P7a
Hsa-Mir-506-P6c
Hsa-Mir-506-P6b
Hsa-Mir-506-P6a2
Hsa-Mir-506-P6a1
Hsa-Mir-506-P5
Hsa-Mir-506-P4b
Hsa-Mir-506-P4a2
Hsa-Mir-506-P4a1
Hsa-Mir-506-P3
Hsa-Mir-506-P29
Hsa-Mir-506-P2
Hsa-Mir-506-P13
Hsa-Mir-506-P12
Hsa-Mir-506-P11
Hsa-Mir-506-P10
Hsa-Mir-506-P1
14.7
0
-6.6
log2(CPM)
adipocyte
artery
bladder
bone
bowel
brain
esophagus
gallbladder
heart
kidney
liver
lung
lymph
muscle
nerve
pancreas
placenta
pleurae
skin
spinal
spleen
stomach
submandibular
testis
thyroid
tongue
trachea
vein
B
Rhesus_MirGeneDB
Mml-Mir-506-P9
Mml-Mir-506-P8
Mml-Mir-506-P7a
Mml-Mir-506-P6c2
Mml-Mir-506-P6c1
Mml-Mir-506-P6b2
Mml-Mir-506-P6b1
Mml-Mir-506-P6a
Mml-Mir-506-P5
Mml-Mir-506-P4f
Mml-Mir-506-P4e
Mml-Mir-506-P4d
Mml-Mir-506-P3
Mml-Mir-506-P29
Mml-Mir-506-P2
Mml-Mir-506-P13
Mml-Mir-506-P12
Mml-Mir-506-P11
Mml-Mir-506-P10
Mml-Mir-506-P1
14.1
0
-3.3
log2(CPM)
bone marrow
brain
heart
kidney
liver
lung
lymph node
skeletal muscle
spleen
testis
thymus
C
Mouse_MirGeneDB
Mmu-Mir-506-P7
Mmu-Mir-506-P27
Mmu-Mir-506-P26
Mmu-Mir-506-P25
Mmu-Mir-506-P24
Mmu-Mir-506-P23
Mmu-Mir-506-P22
Mmu-Mir-506-P21
Mmu-Mir-506-P20
Mmu-Mir-506-P2
Mmu-Mir-506-P19
Mmu-Mir-506-P18
Mmu-Mir-506-P17
Mmu-Mir-506-P16
Mmu-Mir-506-P15d
Mmu-Mir-506-P15c2
Mmu-Mir-506-P15c1
Mmu-Mir-506-P15b2
Mmu-Mir-506-P15b1
Mmu-Mir-506-P15a
Mmu-Mir-506-P14
Mmu-Mir-506-P1
14.5
0
-6.6
log2(CPM)
bone
brain
cerebellum
embryonic
heart
intestine
kidney
liver
lung
muscle
ovaries
pancreas
skin
spinal
spleen
testis
D
Rat_MirGeneDB
Rno-Mir-506-P7
Rno-Mir-506-P6
Rno-Mir-506-P28
Rno-Mir-506-P26
Rno-Mir-506-P25
Rno-Mir-506-P24
Rno-Mir-506-P23
Rno-Mir-506-P22
Rno-Mir-506-P21
Rno-Mir-506-P20
Rno-Mir-506-P2
Rno-Mir-506-P19
Rno-Mir-506-P18
Rno-Mir-506-P17
Rno-Mir-506-P15
Rno-Mir-506-P14
Rno-Mir-506-P1
14.2
0
-5.1
log2(CPM)
adrenal
brain
brainstem
cerebellum
cerebrum
cortex
drg
duodenum
fat subcutaneous
heart
hippocampus
ileum
intestine
jejunum
kidney
liver
lung
medulla
muscle
ovary
pancreas
skin
spleen
stomach
testis
thymus
uterus
whole
E
Rabbit_MirGeneDB
Ocu-Mir-506-P7e
Ocu-Mir-506-P7d
Ocu-Mir-506-P6
Ocu-Mir-506-P5
Ocu-Mir-506-P3
Ocu-Mir-506-P2
Ocu-Mir-506-P1
13.2
0
-2.1
log2(CPM)
brain
heart
kidney
testis
whole
F
Dog_MirGeneDB
Cfa-Mir-506-P7
Cfa-Mir-506-P3b
Cfa-Mir-506-P3a
Cfa-Mir-506-P2e
Cfa-Mir-506-P2d
Cfa-Mir-506-P1
9.3
0
-4.1
log2(CPM)
blood
cerebellum
cortex
heart
hypothalamus
kidney
liver
lung
ovary
skin
testis
G
Cow_MirGeneDB
Bta-Mir-506-P6
Bta-Mir-506-P3
Bta-Mir-506-P2h
Bta-Mir-506-P2g
Bta-Mir-506-P2f
Bta-Mir-506-P1
5.0
0
-2.5
log2(CPM)
adult backfat
bone marrow
brainstem
monocytes
cerebellum
corium
cortex
dental pulp
fetal backfat
heart
hypothalamus
iris
kidney
lactation mammary gland
longissimus thoracis
other mammary gland
optic nerve
oral papillae
penis
retina
lymph node
testis
H
Human_PRJNA686442
4
2
0
−2
−4
FDR
(Testis/Others)
miR-890 7.71E-25
miR-892c 2.67E-26
miR-892a 8.01E-24
miR-892b 1.32E-16
miR-891a 2.63E-25
miR-891b 3.18E-15
miR-888 1.69E-31
miR-513c 1.02E-28
miR-508 1.46E-28
miR-509 1.54E-24
miR-513a 9.86E-30
miR-507 4.25E-22
miR-514a 4.79E-31
miR-506 3.42E-21
miR-514b 3.62E-19
miR-510 1.45E-18
miR-513b 1.02E-28
adipocyte
adrenal−gland
artery
bladder
bone
bowel−colon
bowel−duodenum
bowel−ileum
bowel−jejunum
bowel−small−intestine
brain
brain−arachnoid−mater
brain−cerebellum
brain−dura−mater
brain−frontal−lobe
brain−grey−matter
brain−hippocampus
brain−hypothalamus
brain−medulla−oblongata
brain−nucleus−caudatus
brain−occipital−lobe
brain−parietal−lobe
brain−pituitary−gland
brain−substantia−nigra
brain−temporal−lobe
brain−thalamus
brain−white−matter
diaphragm
esophagus
gallbladder
glandular−breast−tissue
heart
heart−atrial−appendage
heart−atrium
heart−muscle
heart−pericardium
heart−ventricle
kidney
kidney−cortex
limb−muscle
liver
lung
lung−bronchus
lymph−node
mesenteric−adipose
muscle
nerve
pancreas
pleurae
prostate
salivary−gland
sclera
skin
small−intestine
spinal−cord
spleen
stomach
submandibular−gland
testis
thyroid
tongue
trachea
urethra
uterus
vein
I
Rat_PRJNA312384
FDR
(Testis/Others)
miR-201 5.32E-115
miR-3551 4.81E-74
miR-3585 1.25E-110
miR-463 4.82E-52
miR-465 5.71E-56
miR-471 7.07E-41
miR-509 9.86E-71
miR-547 4.79E-57
miR-741 4.72E-50
miR-742 1.48E-34
miR-743a 5.33E-48
miR-743b 7.89E-70
miR-871 4.82E-52
miR-878 1.25E-28
miR-880 2.88E-50
miR-881 8.30E-55
miR-883 2.85E-41
12
4
LogCPM
1 2 3 4 5 6 7 8 9 10 11 12 13 14 15 16 17 18 19 20 21 22 23
Tissues

**Figure S6. sRNA-seq of multiple tissues from different species.** A. sRNA-seq from MirGeneDB across multiple tissues in humans. B. sRNA-seq from MirGeneDB across multiple tissues in rhesus. C. sRNA-seq from MirGeneDB across multiple tissues in mice. D. sRNA-seq from MirGeneDB across multiple tissues in rats. E. sRNA-seq from MirGeneDB across multiple tissues in rabbits. F. sRNA-seq from MirGeneDB across multiple tissues in dogs. G. sRNA-seq from MirGeneDB across multiple tissues in cows. H. sRNA-seq from RPJNA686442 across multiple tissues in humans. The testis sample was labeled in red. FDR was indicated on the right side of the plot. The average value was used for the heatmap, $n \geqslant 2$. I. sRNA-seq from RPJNA312384 across multiple tissues in rats. Testis samples were labeled in red. FDR was indicated on the right side of the plot. Rats' tissues analyzed include: 1. Adrenal, 2. Brainstem, 3. Cerebellum, 4. Cerebrum, 5. Cortex, 6. Dorsal root ganglia, 7. Duodenum, 8. Heart, 9. Hippocampus, 10. Ileum, 11. Jejunum, 12. Kidney, 13. Liver, 14. Medulla, 15. Muscle Biceps, 16. Muscle Soleus, 17. Ovary, 18. Pancreas, 19. Stomach Glandular, 20. Stomach Non-Glandular, 21. Testicle, 22. Uterus, and 23. Whole Blood.

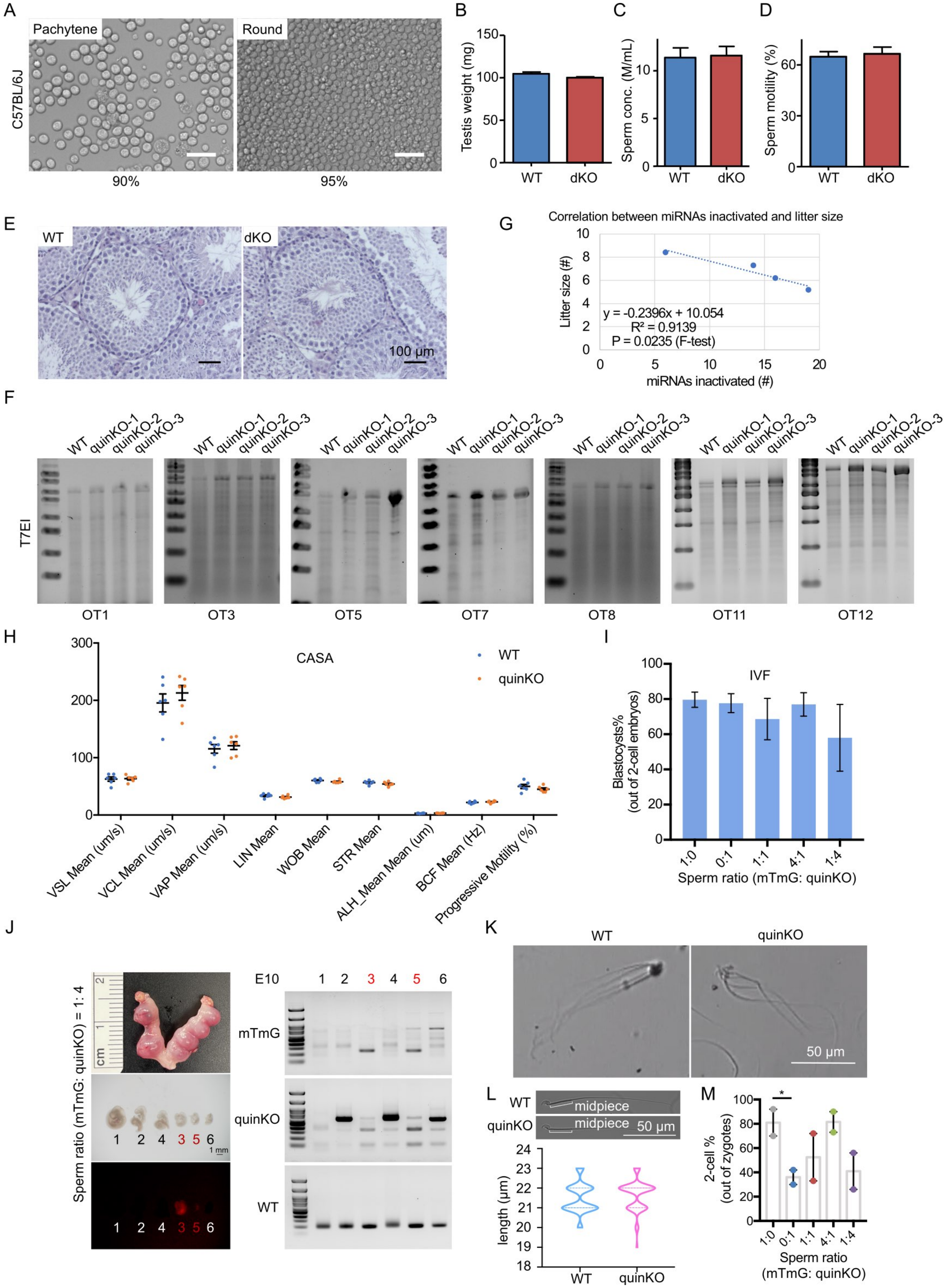
A
Pachytene
Round
C57BL/6J
90%
95%
B
Testis weight (mg)
150
100
50
0
WT
dKO
C
Sperm conc. (M/mL)
10
5
0
WT
dKO
D
Sperm motility (%)
60
30
0
WT
dKO
E
WT
dKO
100 µm
G
Correlation between miRNAs inactivated and litter size
Litter size (#)
10
8
6
4
2
0
y = -0.2396x + 10.054
R² = 0.9139
P = 0.0235 (F-test)
0
5
10
15
20
miRNAs inactivated (#)
F
T7EI
WT quinKO-1 quinKO-2 quinKO-3
OT1
OT3
OT5
OT7
OT8
OT11
OT12
H
300
200
100
0
CASA
WT
quinKO
VSL Mean (um/s)
VCL Mean (um/s)
VAP Mean (um/s)
LIN Mean
WOB Mean
STR Mean
ALH_Mean Mean (um)
BCF Mean (Hz)
Progressive Motility (%)
I
IVF
Blastocysts%
(out of 2-cell embryos)
100
80
60
40
20
0
1:0
0:1
1:1
4:1
1:4
Sperm ratio (mTmG: quinKO)
J
Sperm ratio (mTmG: quinKO) = 1: 4
cm
1
2
1 2 4 3 5 6
1 mm
1 2 4 3 5 6
E10
1 2 3 4 5 6
mTmG
quinKO
WT
K
WT
quinKO
50 µm
L
WT
quinKO
midpiece
midpiece
50 µm
length (µm)
24
23
22
21
20
19
18
WT
quinKO
M
2-cell %
(out of zygotes)
100
80
60
40
20
0
*
1:0
0:1
1:1
4:1
1:4
Sperm ratio
(mTmG: quinKO)

**Fig. S7. Phenotypes of *miR-506* family KO mice.** A. The purity of pachytene spermatocytes and round spermatids after STA-PUT cell sorting in C57BL/6J mice. B. Testis weight of WT and dKO mice. C. Sperm concentrations of WT and dKO males. D. Sperm motility of WT and dKO. E. HE staining of WT and dKO testes. Data presented in panels B, C, and D represent Mean ± SD, n=5. F. T7EI assay on the WT and quinKO mice genomic DNA from tail snips. G. Correlation between miRNAs inactivated and litter size in the 4 KO animal models: *miR-465* sKO, tKO, quadKO, and quinKO. H. Sperm parameters assessed by CASA. n=6 for each sample. VSL, straight-line velocity; VCL, curvilinear velocity; VAP, average path velocity; LIN, linearity; WOB, wobble VAP/VCL; STR, straightness; ALH, amplitude of lateral head displacement; BCF, beat cross frequency. I. Percentage of blastocytes developed from WT MII oocytes fertilized by mixed sperm from mTmG (control) and quinKO males at different ratios. Data were based on three independent IVF experiments. J. Representative embryos and gel images obtained from the co-artificial insemination at embryonic day 10 (E10) using a sperm ratio (mTmG: quinKO) of 1:4. Left panel, a representative image of the embryos collected at E10; right panel, PCR validation of the embryos' genotype. K. Sperm aggregation in WT and quinKO mice. L. Sperm midpiece size of WT and quinKO mice. At least 70 sperm were counted for each sample. Upper panel, representative images of WT and quinKO sperm. M. IVF 2-cell rate of mTmG and quinKO with different sperm ratios. 2-cell ratios were calculated as the number of 2-cell embryos out of the number of zygotes. Data were based on two independent IVF experiments. *, $p < 0.05$, paired t-test was used for the statistical analysis.

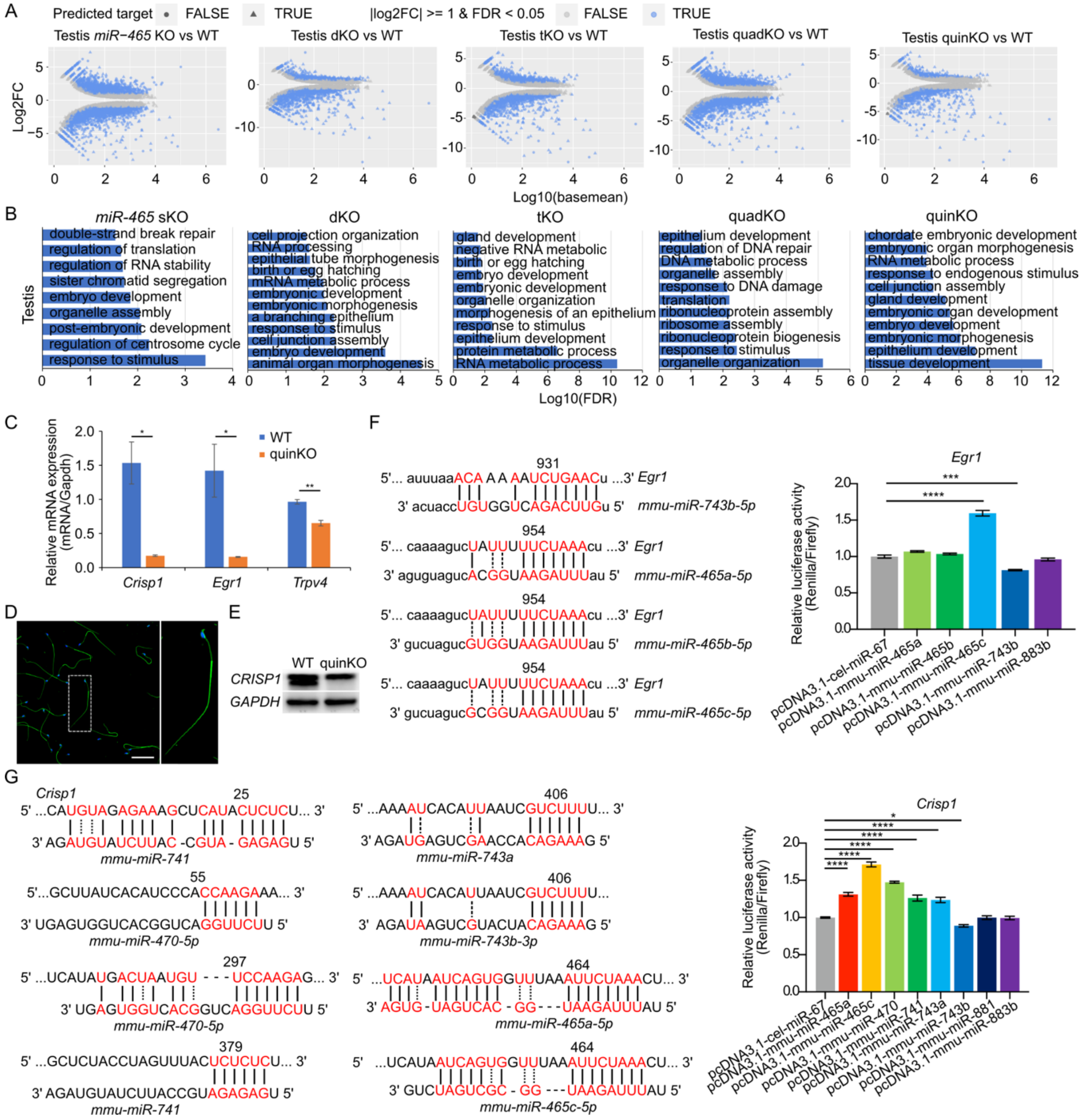


**Figure S8. Dysregulated targets in the X-linked *miR-506* family KO testes.** A. Differentially expressed genes (DEGs) between KO and WT testes, n=3 for each sample. Predicted targets were indicated as the triangle, and the genes with |log2FC| >= 1 and FDR <0.05 were labeled in blue. B. GO term enrichment analyses of the genes fall in differentially expressed (|log2FC| >= 1 and FDR <0.05) and predicted targets, termed as differentially expressed targets (DETs), in KO testes. C. qPCR validation of differentially expressed genes in the quinKO testis. n=3 for each sample. * and ** indicate $p < 0.05$ and 0.01, respectively. T-test was used for the statistical analysis. D. Immunofluorescence of CRISP1 in WT sperm. Scale bar = 50µm. E. Western blot of *CRISP1* in WT and quinKO testis samples. F. Luciferase assay of *Egr1* 3'UTR and *miR-506* family miRNAs from mice. Three biological replicates were done for each sample. Left panel, the predicted binding sites for the *miR-506* family miRNAs and *Egr1* 3'UTR; right panel, luciferase activity of the *miR-506* family miRNAs and the *Egr1* 3'UTR. *** and **** indicate adjusted $p < 0.001$ and 0.0001, respectively. One-way ANOVA was used for the statistical analysis. G. Luciferase assay of *Crisp1* 3'UTR and *miR-506* family miRNAs from mice. Three biological replicates were done for each sample. Left panel, the predicted binding sites for the *miR-506* family miRNAs and *Crisp1* 3'UTR; right panel, luciferase activity of the *miR-506* family miRNAs and the *Crisp1* 3'UTR. *** and **** indicate adjusted $p < 0.001$ and 0.0001, respectively. One-way ANOVA was used for the statistical analysis.

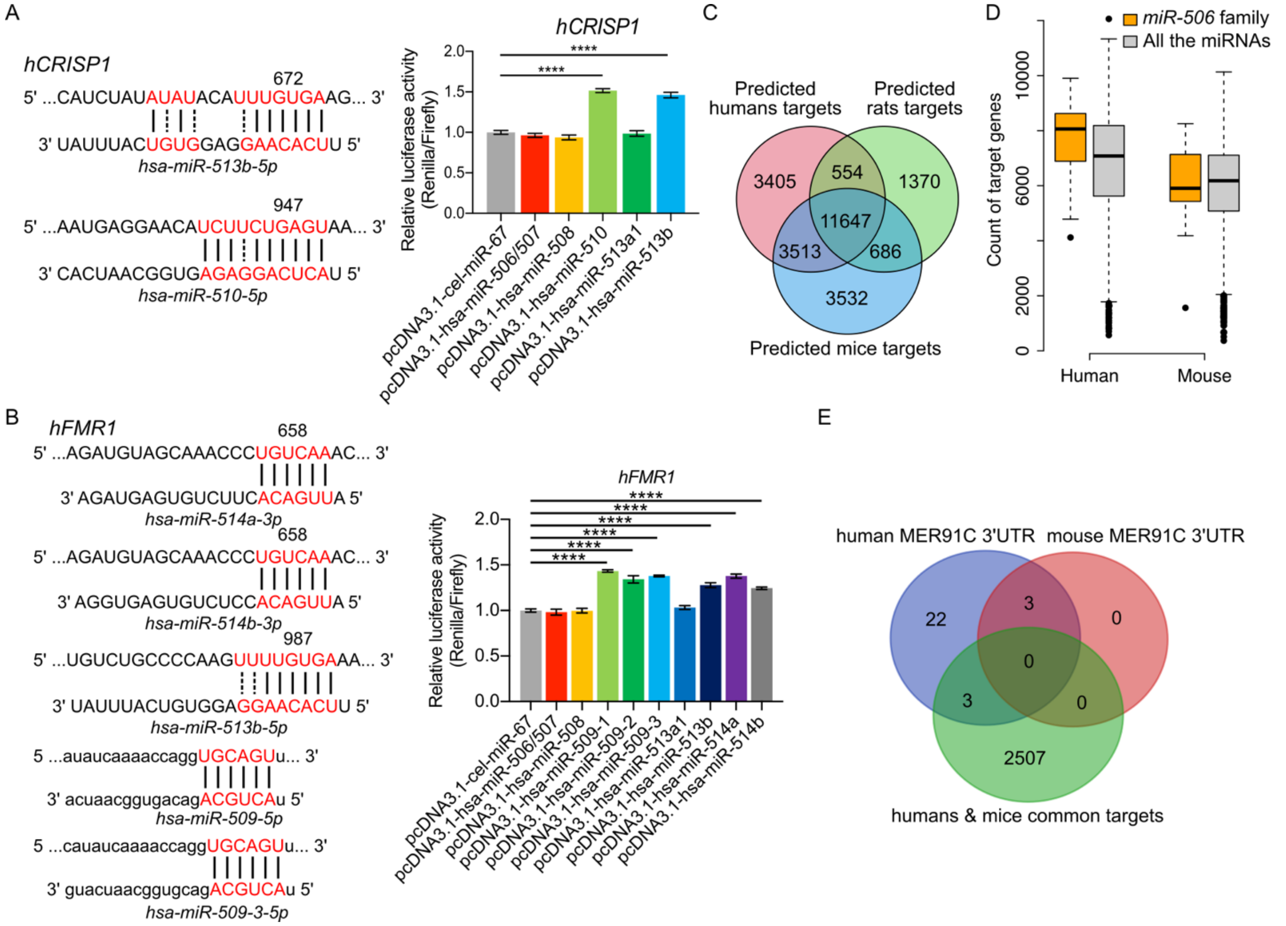


**Figure S9. Dysregulated targets are shared across humans, mice, and rats.** A. Luciferase assay of *CRISP1* 3'UTR and *miR-506* family miRNAs from humans. Three biological replicates were done for each sample. Left panel, the predicted binding sites for the *miR-506* family miRNAs and *CRISP1* 3'UTR; right panel, luciferase activity of the *miR-506* family miRNAs and the *CRISP1* 3'UTR. **** indicates adjusted $p < 0.0001$. One-way ANOVA was used for the statistical analysis. B. Luciferase assay of *FMR1* 3'UTR and *miR-506* family miRNAs from humans. Three biological replicates were done for each sample. Left panel, the predicted binding sites for the *miR-506* family miRNAs and *FMR1* 3'UTR; right panel, luciferase activity of the *miR-506* family miRNAs and the *FMR1* 3'UTR. **** indicates adjusted $p < 0.0001$. One-way ANOVA was used for the statistical analysis. C. Intersections of predicted targets of the *miR-506* family, without using RNA-seq as a reference, in humans, mice, and rats from four databases: miRWalk, TargetScan, miRDB and microrna.org. D. Comparison of the number of the predicted targets, without using RNA-seq as a reference, per miRNA for the X-linked *miR-506* family and all the miRNAs (baseline) in mice and humans. E. Intersections of human and mouse common targets with transcripts containing MER91C DNA transposon in their 3'UTR in humans and mice.
